# Does nonlinear blood-brain barrier transport matter for morphine dosing strategies?

**DOI:** 10.1101/2023.03.17.533135

**Authors:** Berfin Gülave, Divakar Budda, Mohammed AA Saleh, JG Coen van Hasselt, Elizabeth CM de Lange

## Abstract

Morphine blood-brain barrier (BBB) transport is governed by passive diffusion, active efflux and saturable active influx. These processes may be associated with nonlinear concentration-dependencies which impact plasma and brain extracellular fluid (brain_ECF_) pharmacokinetics of morphine. In this study, we aim to evaluate the impact of nonlinear BBB transport on brain_ECF_ pharmacokinetics of morphine and its metabolites for different dosing strategies using a physiologically based pharmacokinetic simulation study. We extended the human physiologically based pharmacokinetic, LeiCNS-PK3.0, model with equations for nonlinear BBB transport of morphine. Simulations for brain_ECF_ pharmacokinetics were performed for various dosing strategies: intravenous (IV), oral immediate (IR) and extended release (ER) with dose range of 0.25-150mg and dosing frequencies of 1-6 times daily. The impact of nonlinear BBB transport on morphine CNS pharmacokinetics was evaluated by quantifying (i) the relative brain_ECF_ to plasma exposure (AUC_u,brainECF_/AUC_u,Plasma_) and (ii) the impact on the peak-to-trough ratio (PTR) of concentration-time profiles in brain_ECF_ and plasma. We found that the relative morphine exposure and PTRs are dose dependent for the evaluated dose range. The highest relative morphine exposure value of 1.4 was found for once daily 0.25mg ER and lowest of 0.1 for 6-daily 150mg IV dosing. At lower doses the PTRs were smaller and increased with increasing dose and stabilized at higher doses independent of dosing frequency. Relative peak concentrations of morphine in relation to its metabolites changed with increasing dose. We conclude that nonlinearity of morphine BBB transport affect the relative brain_ECF_ exposure and the fluctuation of morphine and its metabolites.

**Highlights:**

Nonlinear transport affects relative morphine exposure in brain_ECF_.
Nonlinear transport affects PK fluctuations of morphine in brain_ECF_.
Nonlinear transport affects brain_ECF_ PK relationship of morphine and its metabolites.

**Graphical abstract:** 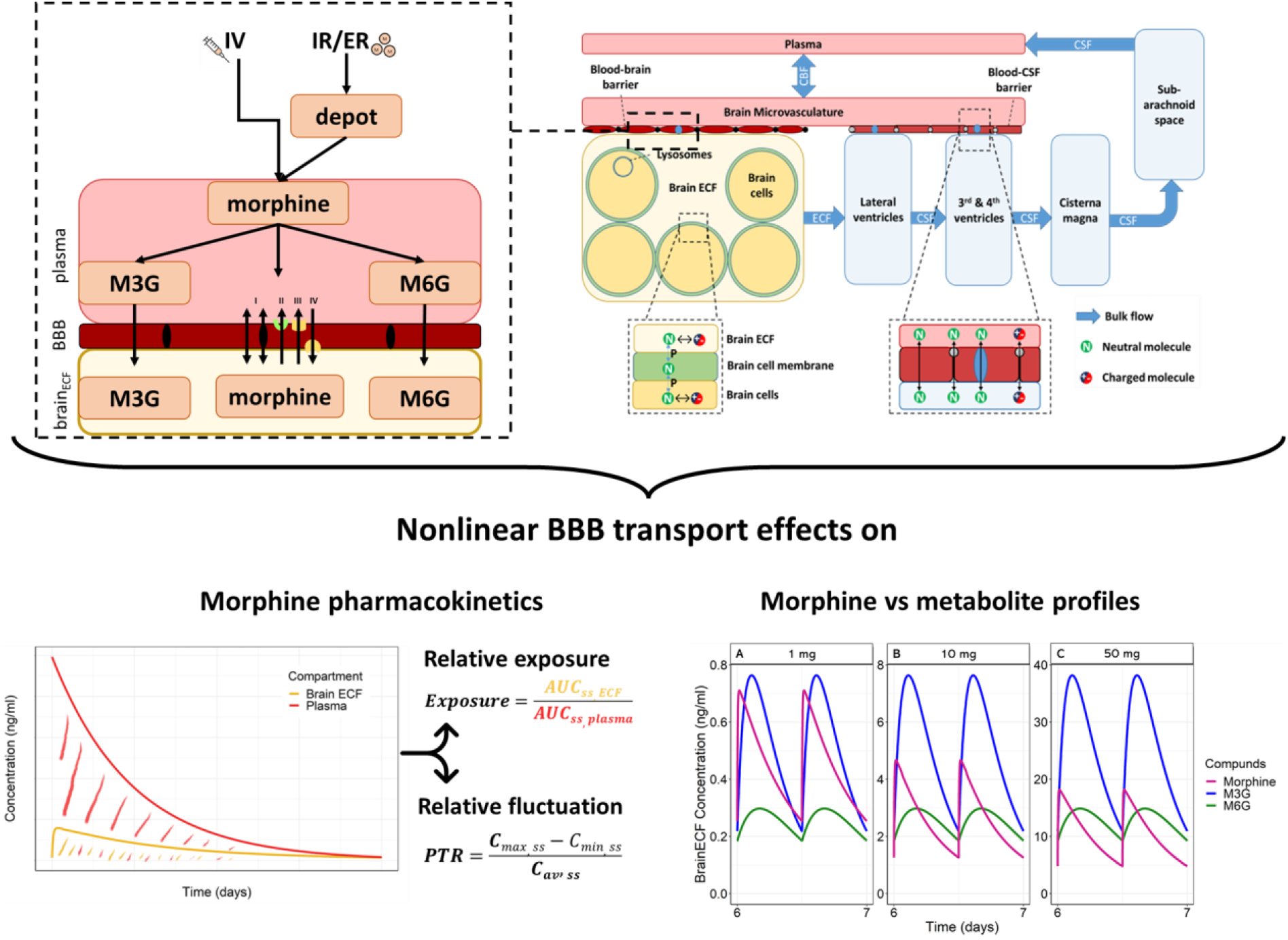

## 1. Introduction

Morphine is an opioid with an important place for the treatment of acute and chronic pain. The main metabolites of morphine in humans are morphine-3-glucuronide (M3G) and morphine-6-glucuronide (M6G) (Christrup, 1997; De Gregori et al., 2012; Frölich et al., 2011). M3G displays a relatively low affinity for opioid receptors and has no analgesic activity. In fact, an opposite effect, hyperalgesia, has been reported (Frölich et al., 2011; Gabel et al., 2022). M6G, however, is capable of eliciting profound analgesic activity, and has even been propose to as the main drive of the analgesic effects of morphine treatment (Klimas and Mikus, 2014; Murthy et al., 2002).

Many pharmacological studies on morphine and its metabolites effect have been performed, but these studies have typically only considered its plasma pharmacokinetics and not the target site pharmacokinetics. However, morphine and its metabolites first need to cross the blood–brain barrier (BBB) to reach the brain extracellular fluid (brain_ECF_) where they can bind with opioid receptors in the brain. Thus, brain_ECF_ concentrations, and not plasma concentrations, should therefore be considered the target site concentration driving the effect. The rate and extent of BBB transport of morphine, M3G, and M6G are different, as has been shown by microdialysis studies in rats. Beside passive transports, para- and transcellular, morphine, M3G and M6G are actively transporter. For morphine, both the P-glycoprotein (P-gp) (Chaves et al., 2017; Letrent et al., 1999; Xie et al., 1999) and probenecid-sensitive transporters (Tunblad et al., 2003) act as BBB efflux transporters, while morphine has a saturable active influx by a yet unidentified BBB influx transporter (Groenendaal et al., 2007; Xie et al., 1999). In rats, it has been shown that blocking P-gp increases the plasma and spinal cord M6G concentrations (Lötsch et al., 2002) while in humans no p-gp related changes in plasma pharmacokinetics were observed (Skarke et al., 2004). The same study in humans showed that probenecid treatment decreases M6G plasma clearance, suggesting that M6G is a substrate for the probenecid-sensitive efflux transporter in the human body (Skarke et al., 2004). It has been reported that GLUT-1 and a digoxin-sensitive transporter can actively efflux M6G but with a weak capacity (Bourasset et al., 2003). For M3G, no P-gp interaction at the level of the BBB has been reported (Xie et al., 1999), while there is a possible involvement of a probenecid-sensitive efflux transporter (Xie et al., 2000). Earlier studies shows the different transport mechanism involved in BBB transport of morphine and its metabolites that can influence their CNS exposure.

When considering BBB transport, constant concentrations at equilibrium (steady-state conditions) and linear pharmacokinetic relationships are often assumed. The ratio from a particular (unbound) concentration in brain and in plasma is used (i.e., a fixed K_p,uu,BBB_ value), without considering (plasma) concentration-dependency (Wright et al., 2011). The concentration-dependency should be considered for drugs with potentially nonlinear active BBB transport processes, and drugs with metabolites that compete in binding to the same receptor(s). Since morphine and its metabolites are known to be affected by nonlinear BBB transport processes, dosing schedules and/or formulations may impact the ultimately observed exposure at the target site.

The aim of this study is to evaluate the impact of nonlinear BBB transport on relative CNS exposure of morphine and its active metabolites for broad range of dosing regimens and formulations. To that end, we will apply a physiologically based pharmacokinetic (PBPK) CNS modeling approach. This PBPK CNS model, the LeiCNS-PK3.0, predicts within two-fold error the unbound drug concentrations at different CNS compartments (Saleh et al., 2021). We expand the LeiCNS-PK3.0 PBPK model with concentration-dependent BBB transport processes of morphine. The area under the curves (AUC) and the peak-to-through ratio (PTR) for unbound plasma and unbound brain_ECF_ pharmacokinetic profiles of morphine and its metabolites were compared to assess the effect of nonlinear BBB transport.

## 2. Methods

### 2.1 Nonlinear transport blood brain barrier

The nonlinear BBB transport of unbound morphine in the LeiCNS-PK3.0 model is described by a concentration-dependent K_p,uu,BBB_ function. To derive this function, a previously published pharmacokinetic model was used that described nonlinear BBB transport of morphine in rats, which included passive diffusion, active efflux and saturable influx transport (Groenendaal et al., 2007). To obtain an equation for K_p,uu,BBB_, this nonlinear model was simulated for rat for a wide range of doses between 0.1 – 500 mg/kg as a continuous infusion for 24 hours to obtain steady-state profiles. We then fitted a power function to relate the plasma unbound steady state concentrations to the K_p,uu,BBB_, resulting in the following power function K_p,uu,BBB_ = 5.4902*C_ss,u,plasma_^-0.552^.

### 2.2 LeiCNS-PK3.0 PBPK model

For this study the previously published CNS PBPK model, LeiCNS-PK3.0 (figure 1B), was used as base model (Saleh et al., 2021). Briefly, this comprehensive model consists of a plasma and multiple CNS and cerebrospinal fluid (CSF) compartments. Between the brain microvasculature and brain_ECF_ and CSF compartments the BBB and the blood-CSF-barrier (BSCFB) are incorporated. The multiple physiological compartments are connected through cerebral blood, brain_ECF_ and CSF flows. Furthermore, this model takes into account pH values in each compartment, as well as brain non-specific tissue binding. As input into the LeiCNS-PK3.0 model-, on one hand, previously published human population plasma pharmacokinetic model for morphine and metabolites following IV and oral dosing was used (figure 1A; table 1) (Oosten et al., 2017). The physicochemical properties of morphine and its metabolites were provided to the model (table 1).

**Figure 1.**
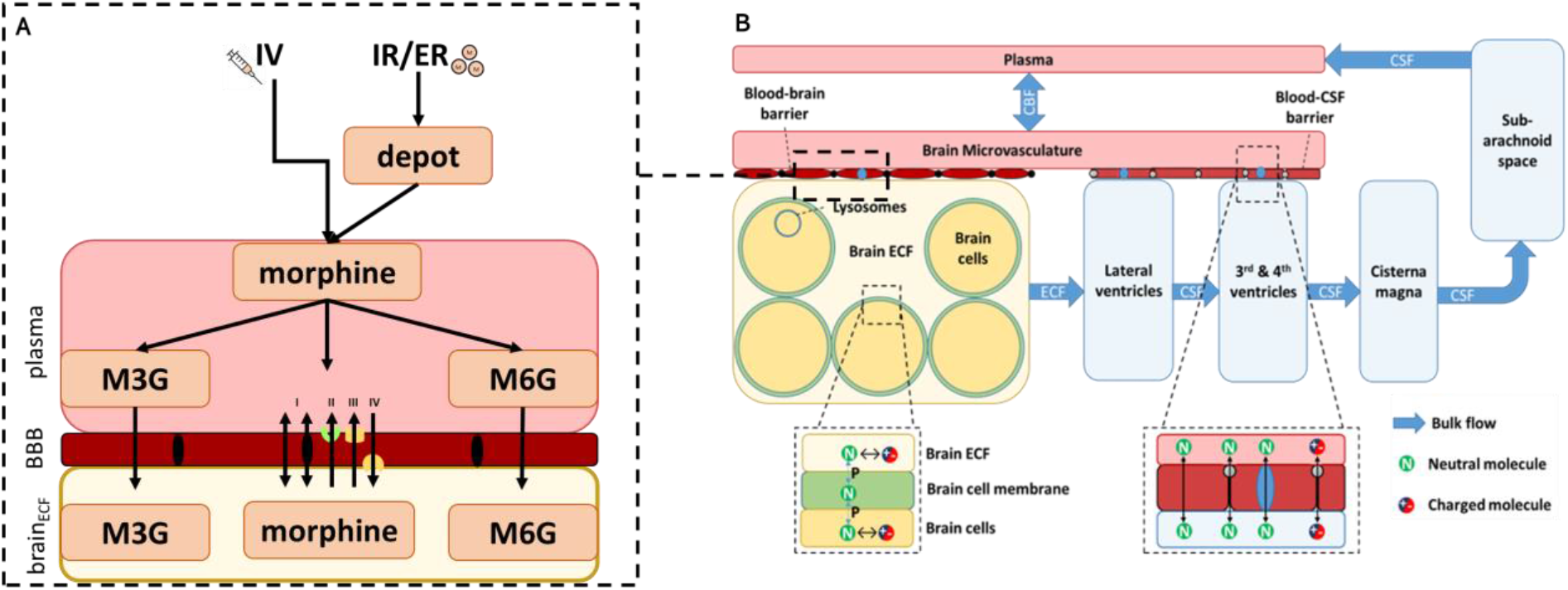
Morphine nonlinear and LeiCNS-PK3.0 model. *A) As plasma input to the LeiCNS-PK3.0 model, previously published plasma pharmacokinetic model by Oosten et al. 2017 was used. This model consisted of one compartment for morphine, morphine-3-glucuronide (M3G) and morphine-6-glucuronide (M6G) in plasma and brain extracellular fluid (brain_ECF_). In this model the nonlinear blood-brain barrier (BBB) transport of morphine by Groenendaal et al. is included while for the metabolites a linear transport across BBB is included. Morphine BBB transport includes I. passive para- and transcellular transport; II. efflux by P-glycoprotein III. efflux by a probenecid sensitive transporter; IV. influx by unidentified saturable influx transporter* (Groenendaal et al., 2007). *B) The LeiCNS-PK3.0 model describes drug distribution in the various CNS compartments, taking into account the drug flow between different physiological compartments, various transport modes, pH influence and non-specific binding. CBF = cerebral blood flow, CNS = central nervous system, CSF= cerebrospinal fluid, ECF = extracellular fluid, P= octanol-water partitioning.*

**Table 1.** Plasma pharmacokinetics and physical-chemical and biological properties of morphine, morphine-3-glucuronide and morphine-6-glucuronide.

|  |  | **morphine** | **M3G** | **M6G** |
| --- | --- | --- | --- | --- |
| **plasma PK** (Oosten et al., 2017) | Central clearance (ml/min) | 1531.67 | 78.5 | 78.5 |
|  | Central compartment volume (ml) | 278000 | 25800 | 25800 |
|  | Absorption rate (min<sup>-1</sup>) | IR: 0.1<br>ER: 0.00368 | - | - |
|  | Oral bioavailability | 0.372 | 0.355<sup>a</sup> | 0.0631<sup>a</sup> |
|  | Fraction formed<sup>b</sup> | 0.323 | 0.573 | 0.104 |
|  | IIV central clearance (as variance) | 0.222 | 0.632 | 0.368 |
|  | IIV central compartment volume (as variance) | 0.747 | 0.247 | 0.243 |
|  | Proportional residual error (as variance) | 0.286 | 0.2 | 0.239 |
| **drug properties** (Wishart et al., 2017) | Molecular weight (g/mol) | 285.34 | 461.46 | 461.46 |
|  | Octanol- water lipophilicity | 0.99 | -0.63 | 0.13 |
|  | Acid ionization constant | 10.26 | 2.67 | 2.87 |
|  | Base ionization constant | 9.12 | 9.17 | 9.12 |
|  | Fraction unbound plasma | 0.65 | - | - |
|  | fAFBBB | 0.21815 | 1 | 1 |
| | $K_{p,uu,BBB}$ | $5.4902 \cdot C_{ss,u,plasma}^{-0.552}$ | 0.1<sup>c</sup> | 0.25<sup>d</sup> |

*a* fraction of metabolite formation in first-pass effect
*b* fraction formed after morphine central clearance
*c* obtained from (Xie et al., 2000)
*d* obtained from (Bouw et al., 2001; Tunblad et al., 2005) *IIV* inter individual variability

To describe physiological processes such as active BBB transport, the model includes asymmetry factors (AF). This value can be seen as the “pure” extent of drug distribution at the barrier, without influences of other elimination routes such as brain_ECF_ bulk flow, which in our model are explicitly separated. The AF are calculated as influx and efflux ratios at steady state and includes K_p,uu,BBB_ values. If a K_p,uu,BBB_ is 1, mainly passive transport is dominating, and the AF_influx_ and AF_efflux_ will be equal. If K_p,uu,BBB_ is lower than 1, AF_efflux_ will be calculated and AF_influx_ set to 1, and the other way around when K_p,uu,BBB_ is higher than 1.

In order to do simulations for humans, a human K_p,uu,BBB_ value is needed as input. The K_p,uu,BBB_ describing morphine nonlinear transport at BBB for human has not been determined. Therefore, the calculated rat K_p,uu,BBB_ power function was used with a translational factor based on a transporter protein expression ratio at human versus rat BBB (abbreviated as fAFBBB) to correct the AF. This factor is only applied when a drug or metabolite is a substrate of a transporter. For morphine, two efflux (P-gp and probenecid-sensitive) and one influx transporter was taken into account. The mean protein expression level of P-gp in humans is 4.21 fmol/μg total protein (Al-Majdoub et al., 2019; Shawahna et al., 2011; Uchida et al., 2011) and in rats this is 19.28 (Al Feteisi et al., 2018; Hoshi et al., 2013), resulting in a ratio of 0.22. For other transporters, such as the probenecid sensitive transporter and the saturable influx transporter, no expression information is available. In this case it was assumed that expression in humans and rats is equal. Since only P-gp is identified an fAFBBB of 0.22 was used for the translation of rat value of AF_BBB_ to that of human. For M3G and M6G no information on the exact transporters is available and for this an fAFBBB of 1 is used.

### 2.3 Simulation scenarios

Morphine and metabolite brain_ECF_ distribution simulations for human were performed for a period of seven days in order to reach a steady state exposure. A dose range of 0.25-150 mg for intravenous (IV), oral immediate release (IR) and extended release (ER) formulations. All the doses are administered once, twice, four and six times a day.

### 2.4 Evaluation of simulation scenarios

To compare the relative morphine exposure, the AUC ratio of brain_ECF_ over plasma AUC at steady-state was used (equation I). The results are also compared for the advised clinical doses for the different formulations.

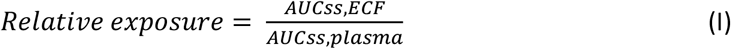

The pharmacokinetic profile fluctuations were evaluated at day seven by comparing the peak-to-trough (PTR) calculated as in equation (II) (Tozer and Rowland, 2016). PTR is calculated by the highest concentration C_max_ minus lowest concentration C_min_ divided by the average concentration C_av_.

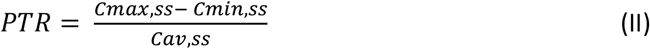

### 2.5 Sensitivity analysis

A sensitivity analysis was performed for the fAFBBB parameter to evaluate the effect of variations of this parameter on the brain_ECF_ exposure (AUC_ECF_). Perturbations of 0.25-2 fold changes in steps of one-quarter of fAFBBB parameter was simulated.

### 2.6 Software

Simulations for nonlinear BBB transport and LeiCNS-PK3.0 models were performed using the package RxODE version 1.1.5 and for sensitivity analysis the additional PKNCA package version 0.9.5 using R version 4.1.3.

## 3. Results

### 3.1 Relative morphine exposure

To compare the effect of nonlinear BBB transport processes, the relative morphine exposure in the brain_ECF_ to plasma was compared for the different formulations at steady state (day seven after treatment start). For all the administration routes, low morphine doses administrations at low frequency resulted in a relative higher exposure of unbound morphine in the brain_ECF_ than in plasma, while increasing dose and frequency led to an increased exposure in plasma compared to brain_ECF_ (figure 2). For almost all administrations, the relative morphine exposure was 1 or lower expect for ER and IR administration of 0.25mg once-a-day. For the metabolites, no differences in relative metabolite exposure have been observed (results not shown). These results show that at low doses (<0.5mg) and low frequency (<twice a day) administrations relative morphine exposure is higher in brain_ECF_ than plasma.

**Figure 2.**
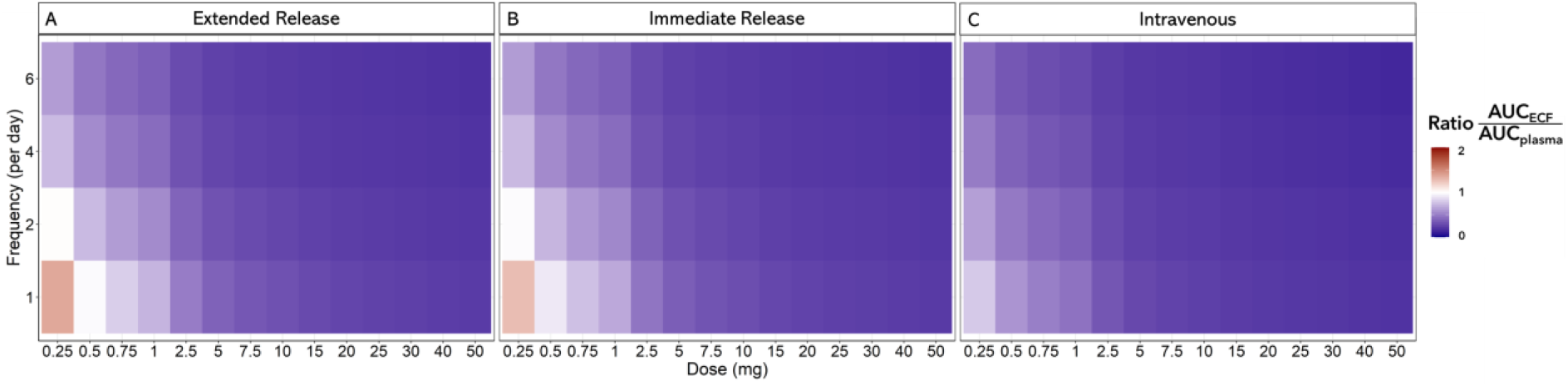
Relative morphine exposure for different formulations and dosing schedules. For A) oral extended release, B) oral immediate release and C) intravenous, the relative morphine exposure is depicted for the different frequencies per day and the dose (and milligrams) administered per time. A ratio of 1 indicates equal exposure of morphine in brain extracellular fluid (ECF) and plasma. Ratio higher than 1 indicates more exposure in brain ECF than plasma while lower than 1 indicates more exposure in plasma than in brain ECF. AUC = area under the curve

### 3.2 Morphine peak-to-trough ratios

To investigate the effect of nonlinear transport on the fluctuation in pharmacokinetic profiles, the PTR concentration ratios for the three dosing regimens were compared. The PTR versus dose in plasma was stable over the simulated dose range, while increasing the frequency, the PTR decreased as expected (figure 3). For brain_ECF_ there was no stable PTR versus dose range observed. The PTR increased with increasing dose and stabilized at higher doses for IV (figure 3), IR and ER (supplementary figure S1 and S2 respectively). The brain_ECF_ PTR over the dosages shows the impact of the saturable influx transporter at lower doses resulting in a nonstable PTR over the dosages. This indicates that steady-state plasma PK profile is not representative for the brain_ECF_ PK profile.

**Figure 3.**
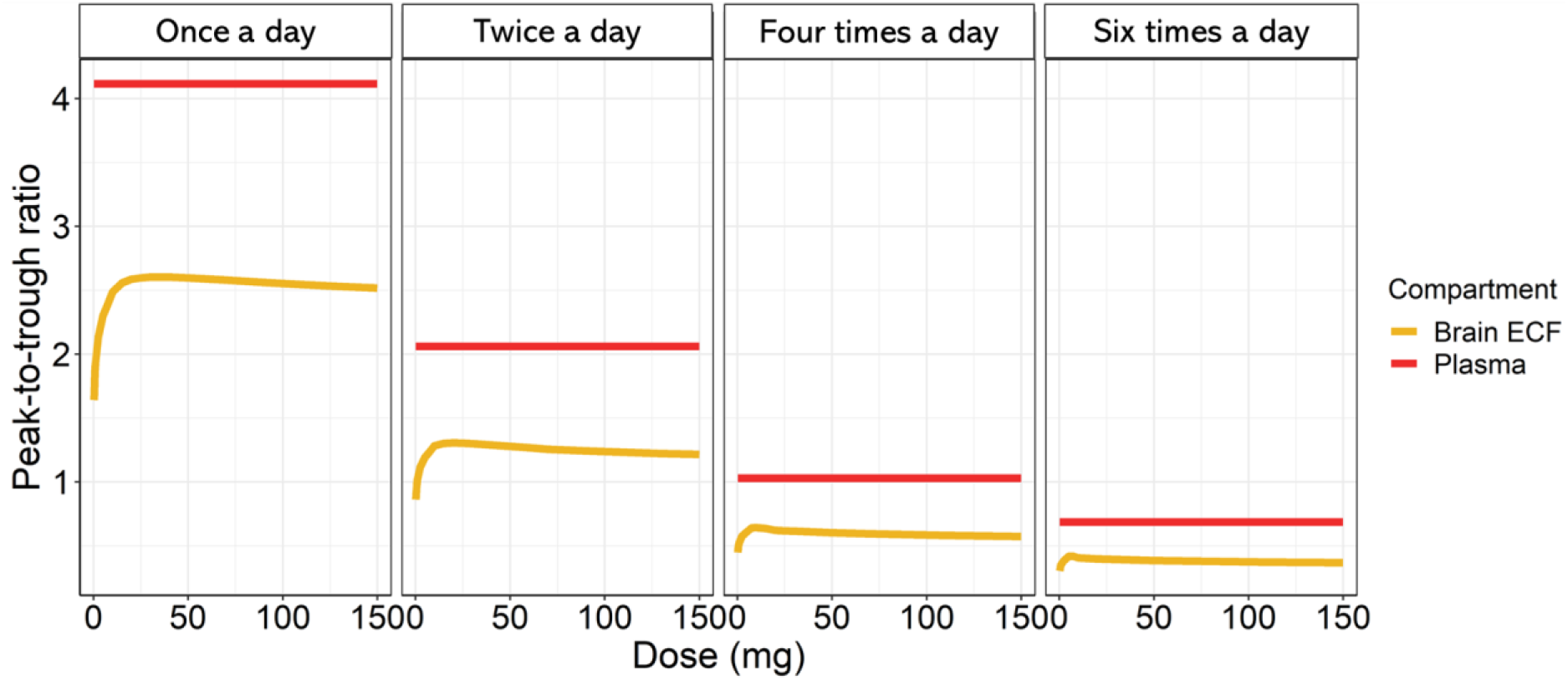
Peak-to-trough ratios (PTR) of unbound morphine in plasma and brain extracellular fluid (brain_ECF_), as a function of the dose. PTR in plasma and brain_ECF_ after IV administration of once, twice, and six times a day.

### 3.3 Nonlinearity effect on metabolite distribution

To study the effect on nonlinear BBB transport of morphine and its metabolites, the profiles are compared for different IV administrations at steady-state. Comparing the peak concentrations (C_max_) of morphine to that of metabolites showed changes with increasing dose. At low dose of 1 mg morphine C_max_ was higher compared to M3G C_max_ and with increasing dose, M3G C_max_ became higher and the difference in peak concentrations of morphine and M3G increased further (figure 4). For morphine versus M6G, this difference in relative C_max_ was other way around, the difference in C_max_ decreased with increasing dose (figure 4). When the metabolite to morphine exposure ratio at steady state was compared (AUCbrain_ECF,metabolite_ / AUCbrain_ECF,morphine_) an increase in this exposure ratio with increasing dose was observed. For M3G/morphine exposure, the ratio at lower doses were above 1 and with increasing dose this ratio increased. Same effect was also observed for M6G/morphine exposure, but at lower dose this ratio was lower than 1 and at higher doses it increased above 1. From these results we can conclude that due to nonlinear BBB transport of morphine, the relation between morphine and metabolites brain_ECF_ peak concentrations and brain_ECF_ exposure ratios changed in relation to dose changes.

**Figure 4.**
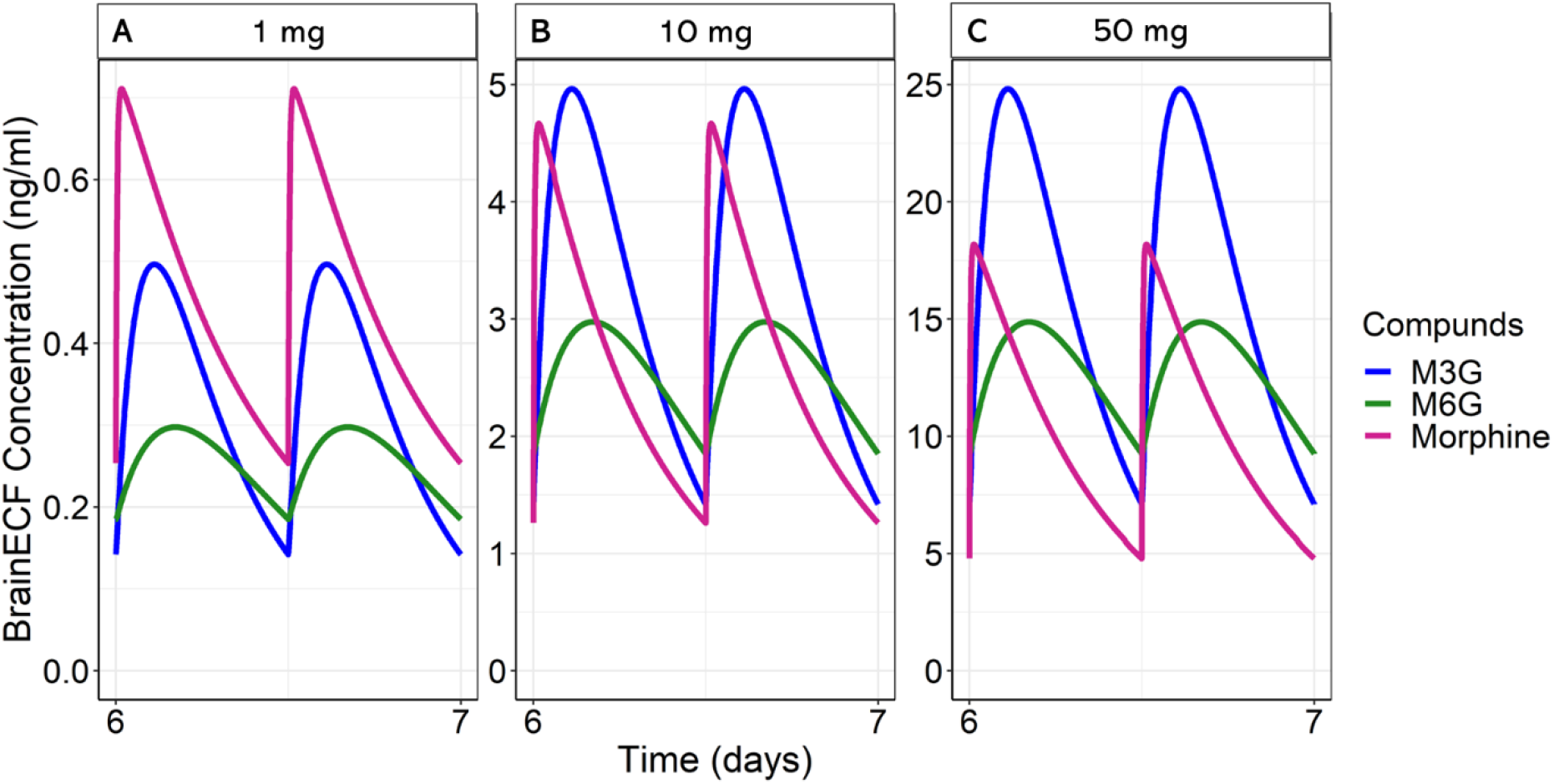
Morphine, morphine-3-glucuronide (M3G) and morphine-6-glucuronide (M6G) distribution in brain extracellular fluid (brain_ECF_) following different dose regimens. The concentration over time profiles of morphine and its metabolites at the brain_ECF_ for 1 (A), 10 (B) and 50 (C) mg twice day IV administration of morphine is compared at steady state (day seven).

### 3.4 Sensitivity analysis

To simulate human morphine brain_ECF_ distribution, rat to human AF_BBB_ factors were translated using the fAFBBB parameter. Morphine is transported by P-gp and one unidentified efflux and one unidentified influx transporter. To calculate fAFBBB to translate rat to human K_p,uu,BBB_ values, the unidentified transporters are assumed to be equally expressed at rat and human BBB. Sensitivity analysis was performed by varying the fAFBBB value to evaluate the possible changes in the brain_ECF_ AUC in case the human BBB transporter expression would deviate from rat values. We find that the highest impact of a change in fAFBBB would be at lower morphine doses, where the contribution of influx transport is the largest (figure 5). For IV administrations of once-a-day, up to 20 mg, an increase in fAFBBB would result in an increase brain_ECF_ AUC. The opposite effect was observed for doses higher than 20mg once a day, where an increase in fAFBBB leading to a decrease in brain_ECF_ AUC. For IR, the possible effect of fAFBBB changes on brain_ECF_ AUC was similar, only the shift in effects occurred at a higher dose of 70mg once a day. For ER, this shift was even at higher dose of 110 mg once a day. The sensitivity analysis showed that the possible largest effect of fAFBBB change would be for a dose of 1 mg within the simulated range of 1 to 150 mg. For ER administrations, a decrease of 75% of fAFBBB would lead to a decrease of 65% brain_ECF_ AUC and an increase of 200% would result in an increase of 74% brain_ECF_ AUC.

**Figure 5.**
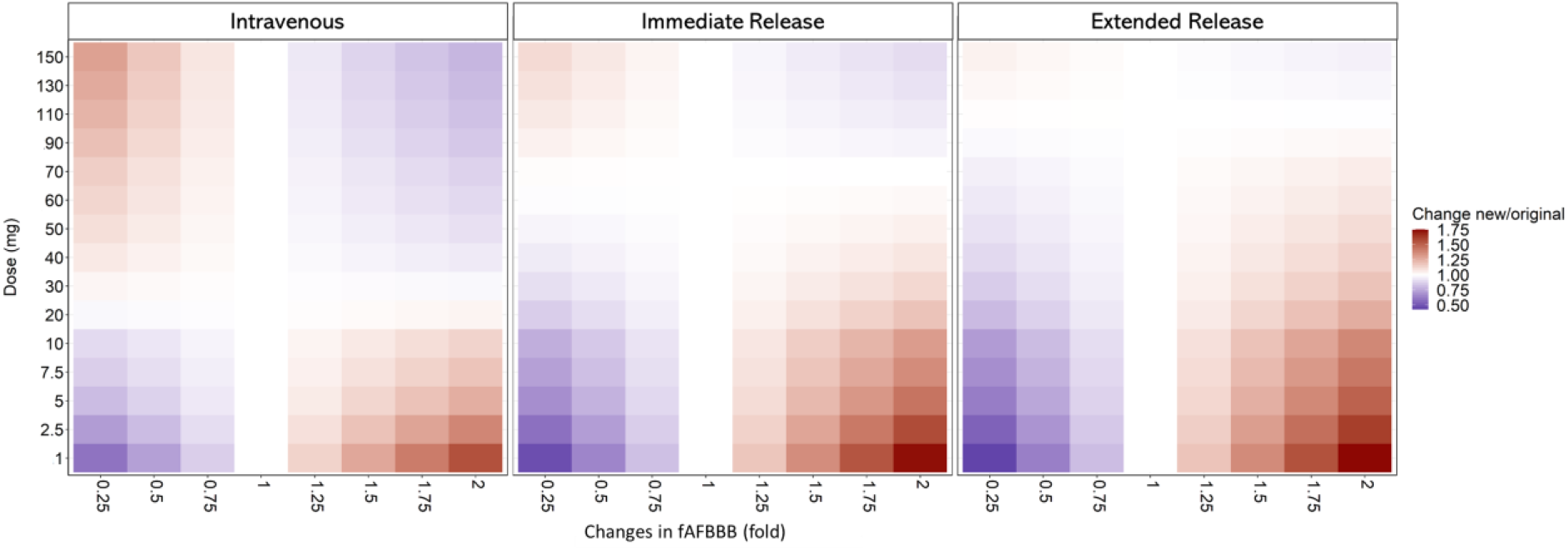
Sensitivity analysis for expression of transporters effect on AUC brain_ECF_) for different dosing regimens. Sensitivity analysis for differences in factor asymmetry factor at blood brain barrier (fAFBBB) and impact on the brain_ECF_ AUC for different doses (1 to 150mg), perturbations (0.25-2-fold changes) and formulations (intravenous, oral immediate and oral extended administrations). Blue indicates a lower new brain_ECF_ AUC compared to original value, white indicates no changes and red a higher new value.

## 4. Discussion

In this study we evaluated the impact of nonlinear BBB transport on distribution of morphine and its active metabolites in the brain_ECF_, by expanding the LeiCNS-PK3.0 PBPK model with nonlinear BBB transport processes. We showed that nonlinear BBB transport of morphine affects the relative target site exposure and PTR, as well as the relation of morphine to its metabolite brain_ECF_ exposure.

Our model predicts the importance of including nonlinear BBB transport to evaluate human brain_ECF_ concentration of morphine and its metabolites. For ethical reasons no such direct information can be obtained from human. In this study, nonlinear BBB transport was implemented for predicting morphine brain_ECF_ pharmacokinetics, based on previous in vivo mice and rat studies that provided quantitative information on plasma concentration dependent BBB influx transport and non-saturable BBB efflux transport processes (Groenendaal et al., 2007; Xie et al., 1999). The extended LeiCNS-PK3.0 model (Saleh et al., 2021; Yamamoto et al., 2017a, 2017b) needs as input a plasma concentration-dependent human K_p,uu,BBB_ (nonlinear BBB transport) of morphine, but such data are not available. So, rat values for concentration-dependent K_p,uu,BBB_ values were derived (Groenendaal et al., 2007), and used in combination with rat to human transporter expression translational factor, fAFBBB, to correct the AF_BBB_. Here the fAFBBB is the relative expression factor of transporters on the BBB in rat and human (Al-Majdoub et al., 2019; Shawahna et al., 2011; Uchida et al., 2011; Yamamoto et al., 2018). Transporters at BBB play a crucial role in drug exposure at brain_ECF_. For this reason, using the relative expression factor as translational factor from rat to human is useful (Yamamoto et al., 2018).

We assumed both unidentified transporters for morphine BBB influx and efflux are equally expressed in rat and human, while the expression of P-gp was scaled from rat to human based on available relative expression values (Al-Majdoub et al., 2019; Shawahna et al., 2011; Uchida et al., 2011; Yamamoto et al., 2018). The sensitivity analysis has shown the possible impact of changes in the fAFBBB on the brain_ECF_ exposure. The results indicate the importance of identification of these transporters, mainly at lower doses. The morphine brain_ECF_ exposure might be mainly at lower doses higher or lower with higher or lower fAFBBB, respectively. For morphine transport across the BBB, P-gp is the only identified active transporter. Another efflux transporter is probenecid dependent as best current knowledge, and furthermore, there is an unidentified saturable influx transporter. Probenecid is known to be an inhibitor for many transporters including multidrug resistance associated proteins (MRPs). The organic anion transporter 1 and 3 (oat1, oat3) and organic anion transporting polypeptide 1 and 2 (oatp1, oatp2) are also inhibited by probenecid (Sugiyama et al., 2001). The possible influx transporter could be the organic cationic transporter 1 (OCT1). Previous study have shown that OCT1 plays a role in hepatocellular saturable and concentration-dependent uptake of morphine (Tzvetkov et al., 2013) and for some cationic compounds in rat and human transfected hepatocytes (Umehara et al., 2007). Whether these suggested transporters are involved in morphine transport their presence at human BBB and transport of morphine should be confirmed.

This study has shown that nonlinear BBB transport mainly affects morphine brain_ECF_ pharmacokinetics at lower dose and lower dosing frequencies for IV, oral IR and ER formulations. With increasing dose, the influx BBB transport of morphine becomes saturated, and its BBB transport becomes mostly linear with plasma concentrations. This nonlinear BBB transport effect is outside the clinical dosing regimens for adult (FDA, 2012, 1984), suggesting no direct impact of nonlinear BBB transport on morphine treatment to adults. For pediatrics, nonlinear BBB transport might have more impact on the treatment regimens. Clinical dosing regimens of morphine in pediatrics depends on their weights resulting for example in oral regimens for pediatrics younger than 12 years a maximum of 200-500 mcg/kg every 4 hours with a maximum of 5mg per day (Unknown author, 2012). With this, the total dose administered of morphine in pediatrics might be within the nonlinear BBB transport dosing range whereby relative more morphine exposure is at the brain_ECF_. Therefore, possible effects due to higher morphine exposure at the brain_ECF_ could be taking into account when administered to pediatrics.

To our best knowledge, nonlinear BBB transport is applicable for morphine, but not for M3G and M6G. The effect of nonlinear BBB transport on the relation of morphine with its metabolites at brain_ECF_ has not been studied before. We found that increasing plasma concentrations result in different brain_ECF_ concentrations ratios of M3G/morphine and M6G/morphine. This may have an impact on their relative receptor binding. The target receptors of morphine and its metabolites are the mu1, mu2, delta and kappa opioid receptors (Imming et al., 2007; Kristensen, 1995). These receptors are predominantly present in the CNS (Peng et al., 2012). For morphine and M6G to exerts their analgesic effect, they should bind to the mu-opioid receptors (Rainville, 2002; Vanderah, 2010; Yamada et al., 2006) and therefore compete with each other. M3G, on the other hand, has a low potency for the mu-opioid receptor (Frölich et al., 2011). In humans, it has been debated that M3G can cause hyperalgesia by binding to Toll-like receptor 4 and may lead to cross-talk of the Toll-like receptor 4 and mu-opioid receptor (Gabel et al., 2022). By binding to the mu-opioid and Toll-like receptors, morphine and M6G actives the Gi-protein and β-arrestin while M3G has a lower potency and activates scaffold proteins (Frölich et al., 2011; Gabel et al., 2022). Altogether, this indicates the need for understanding brain_ECF_ exposure as step one, followed by further exploration on the consequences on receptor binding.

One aspect not yet taken into account in this study is the possible metabolism of morphine within the CNS. This study assumed only metabolism of morphine in the liver and used plasma pharmacokinetics with fixed metabolite fractions (Oosten et al., 2017). However, a previous study has shown possible M6G formation from morphine in human brain homogenates (Yamada et al., 2003), and also in rat microglia (Togna et al., 2013). This is to be further investigated in future research.

## 5. Conclusion

In conclusion, our simulations indicate that nonlinear BBB transport of morphine and its metabolites may affect exposure in brain_ECF_ target site concentrations, in particular at lower doses, enabled by an in silico PBPK modeling approach using the LeiCNS-PK3.0 model.

## 8. Authors contribution

**Berfin Gülave**: Conceptualization, Data collection, preparation and analysis, Model simulations, Writing original draft, review and editing. **Divakar Budda**: Conceptualization, Data collection, preparation and analysis, Model simulations, Writing original draft, review and editing. **Mohammed AA Saleh**: Data preparation and analysis. **JG Coen van Hasselt**: Conceptualization, Writing original draft, review and editing. **Elizabeth CM de Lange:** Conceptualization, Writing original draft, review and editing.

