## Supplementary figures for "Does nonlinear blood-brain barrier transport matter for morphine dosing strategies?"

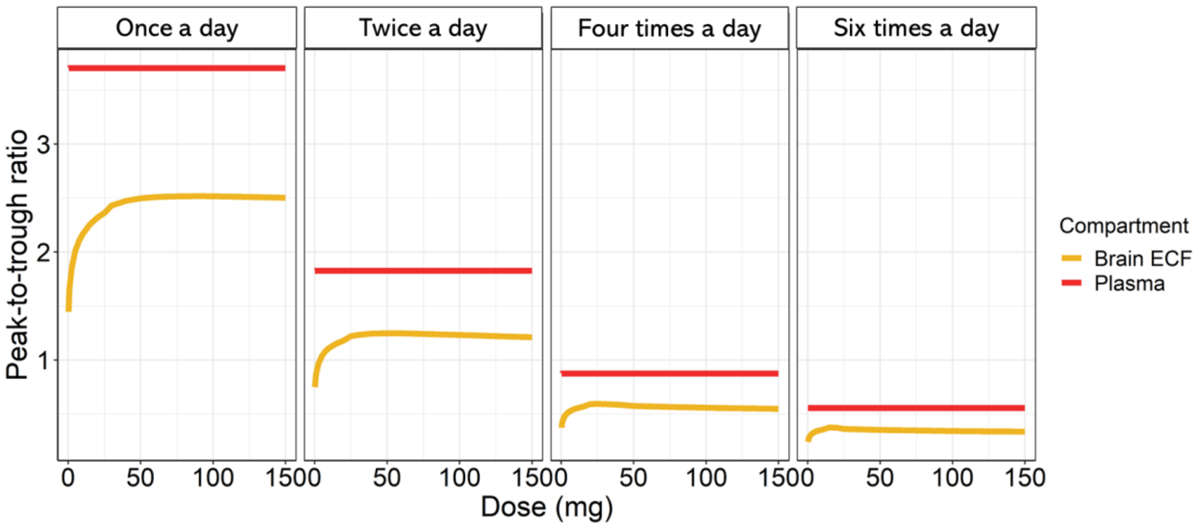


***Figure S1 Peak-to-trough ratios (PTR) of unbound morphine in plasma and brain extracellular fluid (brain$_{ECF}$), as a function of the dose.*** *PTR in plasma and brain$_{ECF}$ after oral immediate release administration of once, twice, and six times a day.*

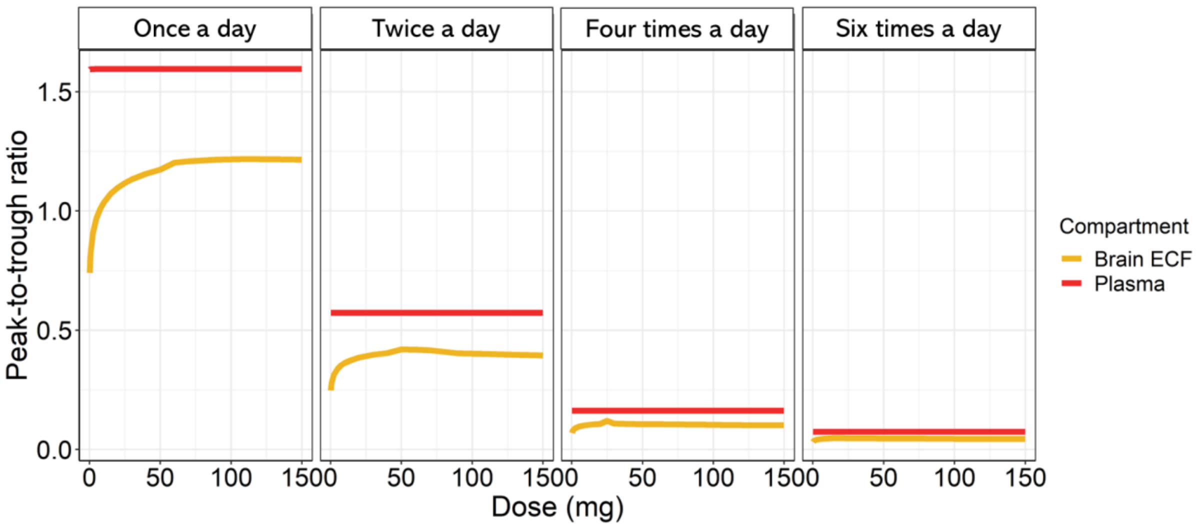


***Figure S2 Peak-to-trough ratios (PTR) of unbound morphine in plasma and brain extracellular fluid (brain$_{ECF}$), as a function of the dose.*** *PTR in plasma and brain$_{ECF}$ after oral extended release administration of once, twice, and six times a day.*
